# A UNIFORM CODING STRUCTURE IN THE NEOCORTEX

**DOI:** 10.64898/2026.05.06.723209

**Authors:** Mohamed Badawy, Alon Amir, Mohammad M Herzallah, Ian T Kim, Drew B Headley, Denis Paré

## Abstract

Prefrontal neurons simultaneously encode multiple task variables. While many studies reported that various groupings of task features could be detected at the population level, the combination of features encoded by individual neurons seemed random. Here, based on unit recordings with Neuropixel probes in behaving rats, we report that far from being random, the representation of information is highly structured. Specifically, the prefrontal network exhibits multiple coding gradients orthogonal to each other in a multidimensional representational space. In this coding structure, neurons have correlated absolute firing rate modulations by different variables, but the polarity of the modulation by one variable is not predictive of that by others. Moreover, this coding structure is manifest in tasks that probe different behavioral processes, ranging from defensive behaviors to sensory discrimination. Last, we find that the same structured representation is apparent in other neocortical regions, including associative and primary sensory areas.

## INTRODUCTION

Neurons in higher-order subcortical and cortical regions like the basolateral amygdala^1–3^, prefrontal cortex^4–7^, and posterior parietal cortex^8–10^ exhibit various forms of multidimensional coding. Such high-dimensional coding schemes are thought to be computationally advantageous as they increase the number of feature combinations that can be discriminated^11^. Although there are regional differences in the type or timing of the variables neurons encode, such as the differential representation of abstract rules in the prefrontal and posterior parietal cortex^12–15^, it was generally reported that coding is strongly task-dependent^14,16–21^. Moreover, many studies commented that the combination of variables encoded by individual neurons defied understanding and seemed random (for instance see ^8,19,22^) as the correlation between the encoding strength of different variables across neurons was close to zero^23^.

Recently, we too generally found low correlations between the coding of different variables by prelimbic neurons. However, we then realized that this result concealed a highly structured representation of information. In this report, we describe this coding structure and show that it is manifest across three tasks that have distinct behavioral and cognitive demands. The variables examined span sensory cues, contextual rules as well as approach and avoidance behaviors emitted spontaneously or in response to conditioned stimuli. Finally, we report that this highly structured representation is also found in other neocortical regions, including associative and primary sensory areas.

## RESULTS

### Tasks

Different rats were trained on one or two of three tasks. Once they became proficient, they were implanted with Neuropixel probes to record unit activity during task performance. The three tasks were (*<u>1</u>*) an operant task that features various conditioned approach and avoidance behaviors called the risk-reward interaction (RRI) task (5 rats), (*<u>2</u>*) a semi-naturalistic foraging task (6 rats), and (*<u>3</u>*) a contextual sensory discrimination task termed the operation task (6 rats).

In the *RRI task*, rats learn that conditioned light stimuli (CSs, 20 s) signal appetitive (CS-R) or aversive (CS-S) outcomes depending on their location (**Fig. 1A1**). When presented behind the west (CS-R1) or east (CS-R2) walls, they indicate that a reward will be delivered there 10 s later. Presentation of the light stimulus under one of three floor sectors (CS-S1, CS-S2, CS-S3) signals that a foot-shock will be delivered there 10 s later. After 7-10 daily training sessions, rats become proficient at this task, successfully earning rewards and avoiding foot-shocks in >80% of trials. See^3^ for a detailed description of behavior in the RRI task.

In the *foraging task*, hungry rats must leave a dimly lit nesting area (**Fig. 1B1**, left) to obtain food pellets located in a brightly illuminated foraging arena (**Fig. 1B1**, right) where they are confronted with a robotic predator on a proportion of trials. Trials start when the doorway to the foraging arena opens. After a delay, rats move to the door threshold and appear to hesitate (waiting phase). Then, they either retreat into the nest or initiate foraging, which typically involves running along a wall, grabbing the food pellet with their mouth, and immediately running back to the nest to consume it. However, when rats approach the predator to obtain food, it surges forward, opening and closing its jaws repeatedly. On predator trials, rats wait longer at the door threshold, and they abort a higher proportion of trials. See^24,25^ for a detailed description of behavior in the foraging task.

**Figure 1.**
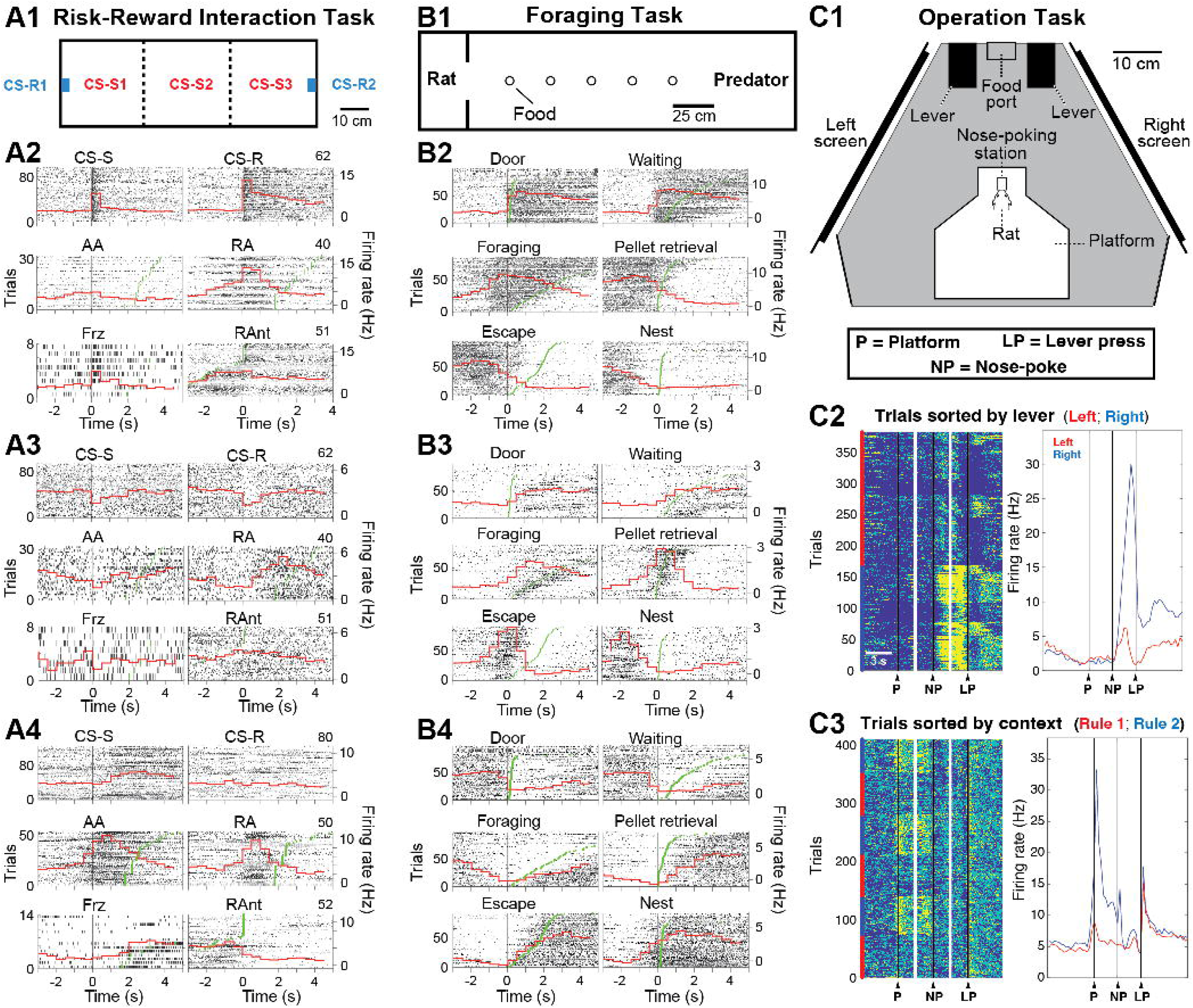
Three behavioral tasks and their neuronal correlates in the prelimbic region. (**A1,B1,C1**) Top view of the behavioral arenas used in the three tasks. (**A2-4, B2-4**) Activity of three different neurons in the RRI (**A2-4**) and foraging (**B2-4**) tasks. In each case, the neuron’s activity is depicted in relation to six task events or behaviors (labels at top) using rasters (each dot or tick stands for one action potential; lines are trials) and PETHs (red) averaging the trials depicted. Green ticks indicate end of the behavior under consideration and trials are rank-ordered by the behavior’s duration (from longest at top to shortest at bottom) except for Reward Anticipation (RAnt) where the green line indicates the end of the preceding behavior (Reward Approach - RA). (**C2,3**) Activity of two prelimbic neurons in the Operation Task. Left, raster where each line is a trial and firing rate is color-coded (blue to yellow indicates low to high firing rates) as a function of real time. Right, PETHs averaging the cells’ firing rates in the trials shown on the left as a function of normalized time. Depending on the panel, trials are sorted by lever (**C2**) or context (Rule 1 or Rule 2). (**C3**). Abbreviations: AA, active avoidance; CS-R, CS predicting a food reward; CS-S, CS predicting a foot-shock; Frz, behavioral freezing; LP, lever press; NP, nose-poke; P, platform; RA, reward approach; RAnt, reward anticipation.

In the *operation task* (**Fig. 1C1**), rats compare visual stimuli shown on two computer monitors placed to their left and right. Two periods of visual stimulation occur on each trial. The first starts when rats step on a platform, triggering the presentation of one of two stimuli identical on both sides (low or high luminance). These stimuli signal the stimulus dimensions rats must heed (target) and ignore (distractor) in the next trial phase (Rule 1 attend to luminance, ignore speed; Rule 2 attend to speed, ignore luminance). The second epoch occurs while rats nose-poke. Here, different stimuli to be discriminated are presented on the left and right: vertically moving circles that vary in speed and luminance, which can assume five values each. Rats must determine on which side the target dimension is highest while ignoring the left-right difference in the distractor dimension. Rats report their decision by pressing a lever on the side where the target dimension is highest. A white light-emitting diode above the reward port signaled correct lever choices. Incorrect lever choices led the two screens to flash. See^26^ for a detailed description of behavior in the operation task.

Below, we will focus on changes in firing rates related to the main task events and behaviors occurring in each of the three tasks. In the *RRI task*, they are the five CSs (CS-R1, CS-R2, CS-S1, CS-S2, CS-S3) and the behaviors they evoked (reward approach and reward anticipation in the case of the CS-Rs; behavioral freezing and active avoidance in the case of the CS-Ss). In the *foraging task*, they are opening of the door (which signals the start of a trial), waiting at the door threshold, foraging, pellet retrieval, escape, and nest re-entry. In the *operation task*, they are the contextual stimuli (Rule 1, Rule 2), the visual stimuli to be discriminated, the left or right lever (through which rats indicate their decision), and whether their decision is correct or incorrect.

### Task-related activity of prelimbic neurons

We recorded prelimbic (PL) neurons while rats performed the RRI, foraging, and operation tasks (534, 577, and 480 single units, respectively). In the three tasks, most cells showed significant changes in firing rates in relation to one or more task events or behaviors, here defined as a deviation from the pre-event baseline beyond *p*<0.05 according to two-tailed signed-rank tests Bonferroni-corrected for the number of task events and behaviors considered in each task (RRI task, 387 of 534 or 72.5%; foraging task, 471 of 577 or 81.6%; operation task, 436 of 480 or 90.8%). The lower part of **figure 1A-C** shows example rasters and peri-event time histograms (PETHs) for a few of the many combinations of significant task-related changes in firing rates observed in the three tasks.

To examine task-related changes in firing rates on a common scale, we computed modulation indices whereby the firing rate change observed in response to a given task event was normalized to the neuron’s baseline firing rate (see Methods). We then plotted the normalized changes in firing rates associated with different pairs of variables. If PL neurons encode a random combination of features, we would expect such scatterplots to exhibit circular or oval distributions centered on the origin. By contrast, in the three tasks, the resulting scatterplots consistently had a distinctive X-shaped profile (**Fig. 2A-O**). This pattern was due to the presence of two diagonals, each reflecting a continuous distribution of neurons with positively vs. negatively correlated changes in firing rates in response to the task events. As shown in **figure 2P-R**, separately averaging the event-related activity of all neurons in each quadrant revealed response patterns that matched the modulation indices of the variables considered. The overall Pearson *r* in each scatterplot varied depending on the balance between the number of neurons in the two diagonals, from as high as 0.79 when most neurons were in the main diagonal (**Fig. 2A-C,K,L**) to near zero when neurons were distributed more evenly in the two diagonals (**Fig. 2H-J,N,O**).

**Figure 2.**
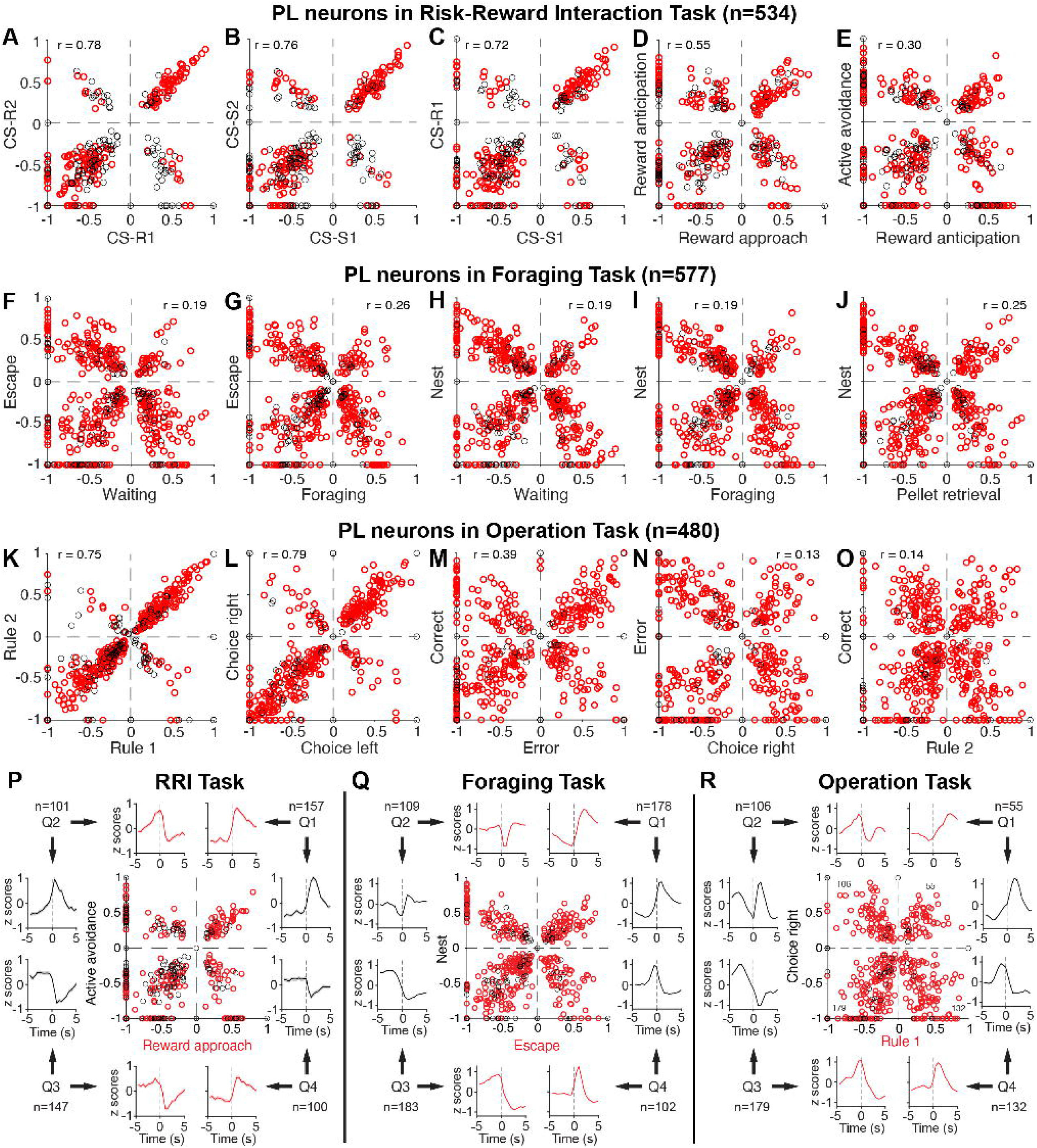
Correlations between the normalized firing rate changes associated with different task events or behaviors reveal a common coding structure across tasks. (**A-E**) RRI task. (**F-J**) Foraging task. (**K-O**) Operation task. Each circle is a prelimbic neuron. *Red circles*: neurons with significant changes in firing rates in relation to at least one of the two task events or behaviors. *Black circles*: cells whose firing rate changes did not reach significance for either. Cells at 0 had no change in firing rate in relation to that variable. The firing rate of the cells at -1 dropped to zero during the response period. The overall Pearson correlation across all four quadrants is indicated in each panel. (**P-R**) *Center:* scatterplot relating the firing rate changes associated with (**P**) active avoidance (y-axis) and reward approach (x-axis) in the RRI task, (**Q**) with nest re-entry and escape in the foraging task, and (**R**) with choice right and rule 1 in the operation task. The Q in Q1-Q4 stands for quadrant. *Periphery:* PETHs of neuronal discharges ± SEM (shading) computed around the onset of a task event or behavior (dashed lines, zero time). PETHs were computed for neurons in each quadrant separately. The number of neurons in each quadrant is indicated. See also **figure S1.**

In the three tasks, there was a negative correlation between firing rates and absolute normalized modulations by all task events (foraging task, *r* = –0.59, *p* = 3.4e-229; RRI task, *r* = – 0.46, *p* = 3.9e-128; operation task, *r* = –0.38, *p* = 1.3e-92; examples in **Fig. S1A**), with presumed fast-spiking interneurons, identified by their short (<0.5 ms) spike waveforms^27^, generally showing low modulations (**Fig. S1B**). These observations raised the possibility that the X-pattern was an artifact of the normalization procedure, causing cells with the lowest firing rates to have the strongest modulations, thereby artificially magnifying the X-pattern. To examine this possibility, we excluded cells with firing rates < 1.8 Hz and “thinned” the spike trains of the remaining neurons to an average firing rate of ∼2 Hz (see Methods). We performed this control analysis in the two tasks with the highest number of available trials, namely the foraging and operation tasks.

Indicating that the X-pattern was not an artefact of the normalization procedure, it persisted after thinning (**Fig. S1C**). This was quantified in two ways. First, we computed the Euclidian distance of neurons from the scatterplots’ origin and compared the results obtained in neurons with thinned vs. not thinned spike trains. Averaging across all pairs of variables, we found that the Euclidian distance differed slightly (by less than 11%) and significantly in both tasks, but in opposite directions (Foraging task: thinned 0.590±0.003, not thinned 0.529±0.003; Operation task, thinned 0.576±0.003, not thinned 0.601±0.003; rank-sum tests, *p’s* ≤ 8.1e-8). Second, for all combination of task variables, we compared the correlation between the absolute modulations in each quadrant separately, again excluding cells with firing rates <1.8 Hz. Averaging across all pairs of variables, we found that correlations in neurons with thinned spike trains remained significant (p<0.01) although they were somewhat lower than in the non-thinned sample (Foraging task: thinned 0.42±0.03, not thinned, 0.59±0.02, rank-sum test, *p* = 6.3e-5; Operation task, thinned 0.43±0.03, not thinned 0.47±0.03; rank-sum tests, *p* = 0.22), which was to be expected given the random removal of spikes in the thinned sample. Taken altogether, these analyses indicate that the X-pattern was not an artefact of the normalization procedure.

### Coding structure in the prelimbic cortex

However, the above analyses are based on PETHs, which cannot readily isolate the firing modulations associated with concurrent events such as the CSs, the motivated behaviors they elicit, and the non-specific influence of locomotion. In the RRI task for instance, at variable delays after CS-R onset, rats approach the reward port and then exhibit reward anticipation by placing their forelimbs on the reward port. Similarly, the CS-Ss elicit different defensive behaviors such as behavioral freezing and/or active avoidance with large variations in relative timing and duration between trials. To circumvent the limitations of PETHs, we fit the firing of neurons with group Lasso generalized linear models (GLMs) with ten-fold cross validation. This approach relies on between-trials variations in the timing and duration of different variables, allowing the model to disambiguate which one(s) neurons encode. Moreover, these GLMs promote sparsity when ascertaining which variables neurons encode and penalize highly correlated predictors ^28–30^.

Although the GLMs used in the three tasks differed (see Methods), they shared the following features. First, they assumed that spiking related to externally-driven events like the CSs in the RRI task are time-locked to the event’s onset. Second, they assumed that spiking related to behaviors anticipate the start of the behavior and continue until its end. Comparisons between the GLMs’ output and PETHs referenced to different variables established the face validity of this approach (**Figs. S2, S3**). At the population level, the average Pearson correlation between the actual and GLM-predicted firing across the variables of interest was 0.676±0.003 in the RRI task (mode: 0.88), 0.698±0.004 in the foraging task (mode: 0.96), and 0.763±0.004 (mode: 0.96) in the operation task.

In the RRI task (**Fig. 3A-J**), GLM-estimated firing modulations associated with different CSs were strongly correlated (*r* ≥ 0.71, *p* ≤ 1e-6), whether the two CSs predicted the same (**Fig. 3A,B**) or different (**Fig. 3C**) outcomes. We interpret this representational alignment as evidence that most PL neurons respond to the CS-Rs and CS-Ss based on their arousing properties, not according to valence. This contrasts with recent models that emphasize a more separable geometric representation of salience and valence in PL^31^. However, despite being dominated by this shared component, the population activity did not collapse into a one-dimensional axis. Instead, ∼90° from the dominant diagonal was a smaller subset of neurons whose modulation by the same variables was inversely correlated, again making the scatterplot look like an X.

**Figure 3.**
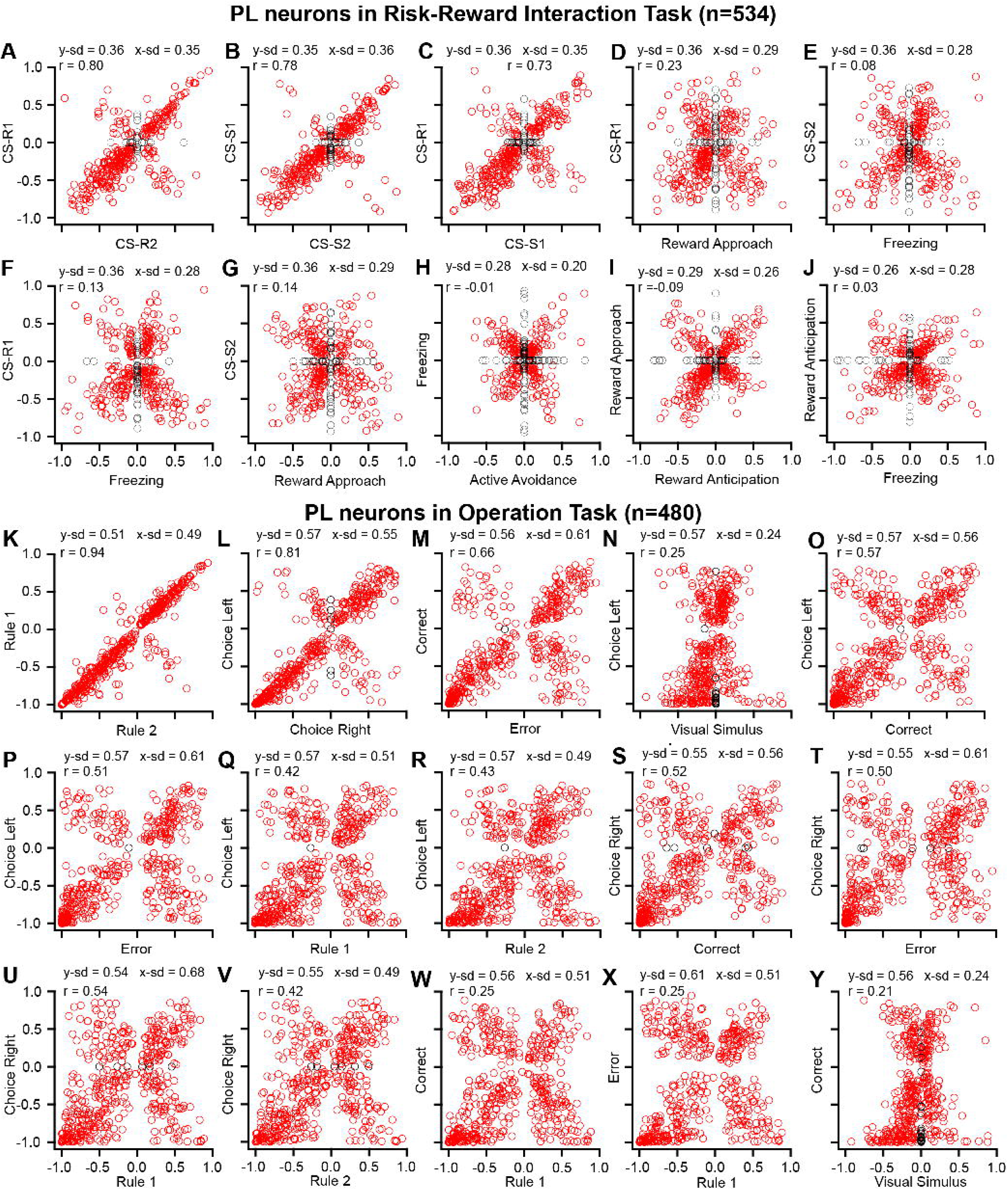
Coding structure in the RRI and operation tasks. Correlation between the GLM-estimated firing modulations associated with different variables of the RRI task (**A-J**) and operation tasks (**K-Y**). Each circle is a prelimbic neuron. *Black circles:* neurons with zero modulation for one or both variables. *Red circles:* neurons with non-zero modulations. At the top of each scatterplot are, from left to right, the Pearson correlation between the variables (*r*), the SD of the variable plotted long the y-axis (y-sd) and of the variable plotted along the x-axis (x-sd). See also **figures S2-S5.**

This X-pattern was also observed when we correlated the firing modulations associated with the CSs and the behaviors they evoked (**Fig. 3D,E**), the CSs and oppositely valenced behaviors (**Fig. 3F,G**), and behaviors of the same (**Fig. 3H,I**) or opposite valence (**Fig. 3J**). In those cases however (**Fig. 3D-J**), the number of cells in the two diagonals was more balanced than in scatterplots relating two CSs. Specifically, the ratio of neuron numbers in the minor to dominant diagonal was 0.19±0.03 when correlating modulations related to two CSs (10 variable pairs) compared to 0.79±0.03 when correlating modulations linked to a CS and a behavior (20 variable pairs; rank-sum test, *p* = 2.81e-7). This indicates that the relative contribution of each diagonal changes depending on the variables compared. Also, note that the angles between the two diagonals depended on the similarity between the standard deviation (SD) of the modulations considered (labels at the top of each panel), with similar or different SDs resulting in equal (**Fig. 3A**) or dissimilar (**Fig. 3F,Y**) adjacent angles, respectively.

The same pattern of results was obtained in the foraging (**Fig. S4**) and operation tasks (**Fig. 3K-Y**). In the latter, scatterplots of the firing modulations associated with the rule stimuli (**Fig. 3K**) bore a striking resemblance to those of the CSs in the RRI task (**Fig. 3A-C**). That is, while modulations linked to these rule stimuli were strongly correlated (*r* = 0.94, *p* < 1e-5), a small subset of inversely correlated neurons was apparent ∼90° from the dominant diagonal. Also paralleling the RRI task, the number of neurons in the two diagonals was less balanced when correlating the modulations linked to the two rule stimuli than when correlating modulations associated with a rule stimulus vs. other variables (ratio of neurons in the minor to dominant diagonals of 0.06 vs. 0.57±0.04, respectively). Last, the relation between the SDs of the modulations and the similarity between adjacent angles in the X-pattern was also manifest in the operation task. Indeed, all scatterplots including modulations associated with the visual stimuli to be discriminated (**Fig. 3N,Y**) had dissimilar adjacent angles because visual stimuli generally elicited little to no response in PL neurons, resulting in a lower SD than with all other variables.

### Coding structure in other neocortical areas

To determine whether the coding structure evidenced above is specific to PL, we examined the activity of V1/V2 neurons (n=694) in the operation task. This analysis revealed that the X-pattern was even more marked in V1/V2 (**Fig. 4**) than in PL (**Fig. 3K-Y**). This visual impression was confirmed by correlating the absolute modulations of all variable pairs (n=21) in the individual quadrants. Absolute correlations were significantly higher in V1/V2 (0.64±0.01) than PL neurons (0.52±0.02; signed-rank test, *p* = 4.21e-10). Other notable differences between the two regions included the expected fact that the visual responses were larger in V1/V2 than PL neurons (absolute modulation by the visual stimulus: V1/V2=0.350±0.008; PL=0.195±0.008; rank-sum test, *p* < 0.0001). Also, in the scatterplot relating the modulations linked to the rule stimuli, the proportion of V1/V2 neurons in the two diagonals was more similar (**Fig. 4A**) than in PL (**Fig. 3K**; ratio of neurons in the minor to dominant diagonals: V1/V2=0.39; PL=0.06).

**Figure 4.**
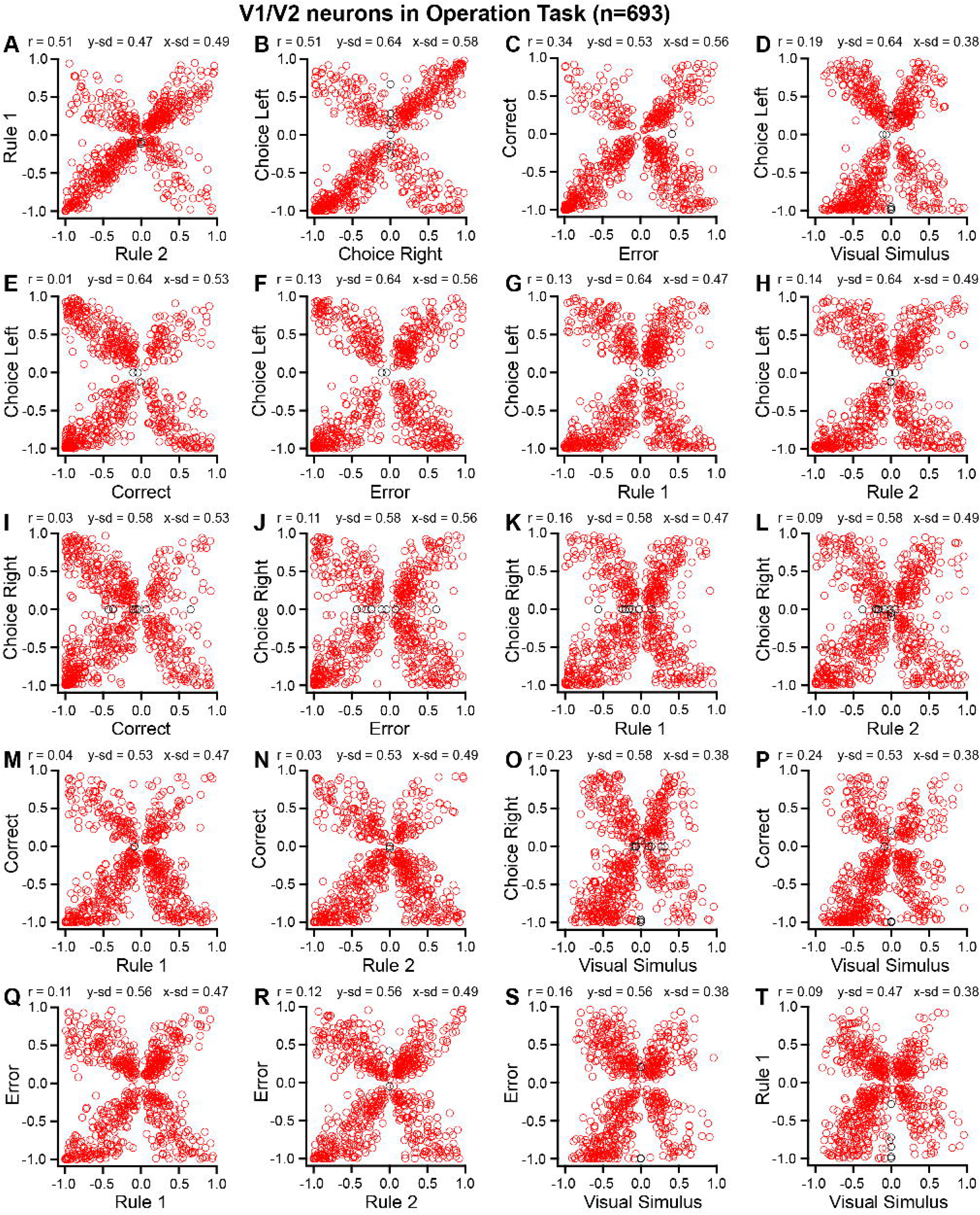
V1/V2 neurons show a similar coding structure as prelimbic cells in the operation task. Correlation between the GLM-estimated firing modulations associated with different variables of the operation task. Each circle is a V1/V2 neuron. *Black circles:* neurons with zero modulation for one or both variables. *Red circles:* neurons with non-zero modulations. At the top of each scatterplot are, from left to right, the Pearson correlation (*r*) between the variables, the SD of the variable plotted long the y-axis (y-sd) and of the variable plotted along the x-axis (x-sd). (**A-C**) Firing modulations estimated in distinct subsets of trials: rule 1 vs. 2 (**A**), left vs. right choice (**B**), and correct vs. error trials (**C**). Various combinations of other variables are shown in panels **D-T**. See also **figures S3 and S5.**

To test whether the X-pattern could arise by chance, we shuffled the modulations of each neuron 1000 times, each time computing the correlations in the four quadrants separately for each pair of variables to obtain null distributions. Indicating that the X-pattern is not a random occurrence, in all three tasks and types of neurons (PL or V1/V2), the actual correlations exceeded 99.99% or more of the shuffled ones (**Fig. S5**). When restricting the analysis to neurons with firing rates >1.8 Hz and spike trains thinned to ∼2 Hz, the actual correlations exceeded 99.4% or more of the shuffled ones.

Finally, given the similarities between results obtained across cortical regions and tasks, readers may wonder whether the X-pattern is ubiquitous in the neocortex. Supporting this possibility, it was also robustly expressed by samples of posterior parietal neurons (n=378) recorded during the operation task as well as S1 neurons recorded during the RRI (n=273) and foraging (n=383) tasks (**Fig. S6**).

### Relation between coding in different tasks

An interesting question is whether the coding structure described above extends to the variables of different tasks. To test this possibility, we took advantage of the fact that 401 PL neurons were recorded in the RRI and foraging tasks on the same day. As shown in **figure 5**, the X-pattern was markedly reduced when relating modulations associated with the variables of different tasks. This was quantified by comparing the absolute correlations of the modulations in the individual quadrants of all variable pairs across tasks vs. within tasks. A Kruskal Wallis ANOVA revealed a significant effect of condition (*df* = 2, H = 116.63, *p* < 0.0001). Post-hoc Dunn tests confirmed that the absolute correlation of the modulations in the individual quadrants was lower across tasks (0.31±0.01) than within the foraging (0.52±0.01, *z* = 9.018.36, *p* < 0.0001) or RRI task (0.49±0.02, *z* = 8.36, *p* < 0.0001), with no difference between the latter two tasks (*z* = 1.69, *p* = 0.09). Suggesting that the attenuation of the X-pattern did not result from poor tracking of single units between tasks, average firing rates in the two tasks were strongly correlated (**Fig. 5I**), more so than expected by chance (**Fig. 5J**). Moreover, a similarly high correlation was found between spike durations (**Fig. 5K**) and spike waveforms (**Fig. 5L**) in the two tasks. The near-zero correlations between functionally analogous variables across tasks, such as escape and active avoidance (r = -0.07), and between cues such as door opening and CS-S1 or CS-R1 (r = 0.16 and 0.07, respectively) indicate that the coding structure does not generalize across tasks.

**Figure 5.**
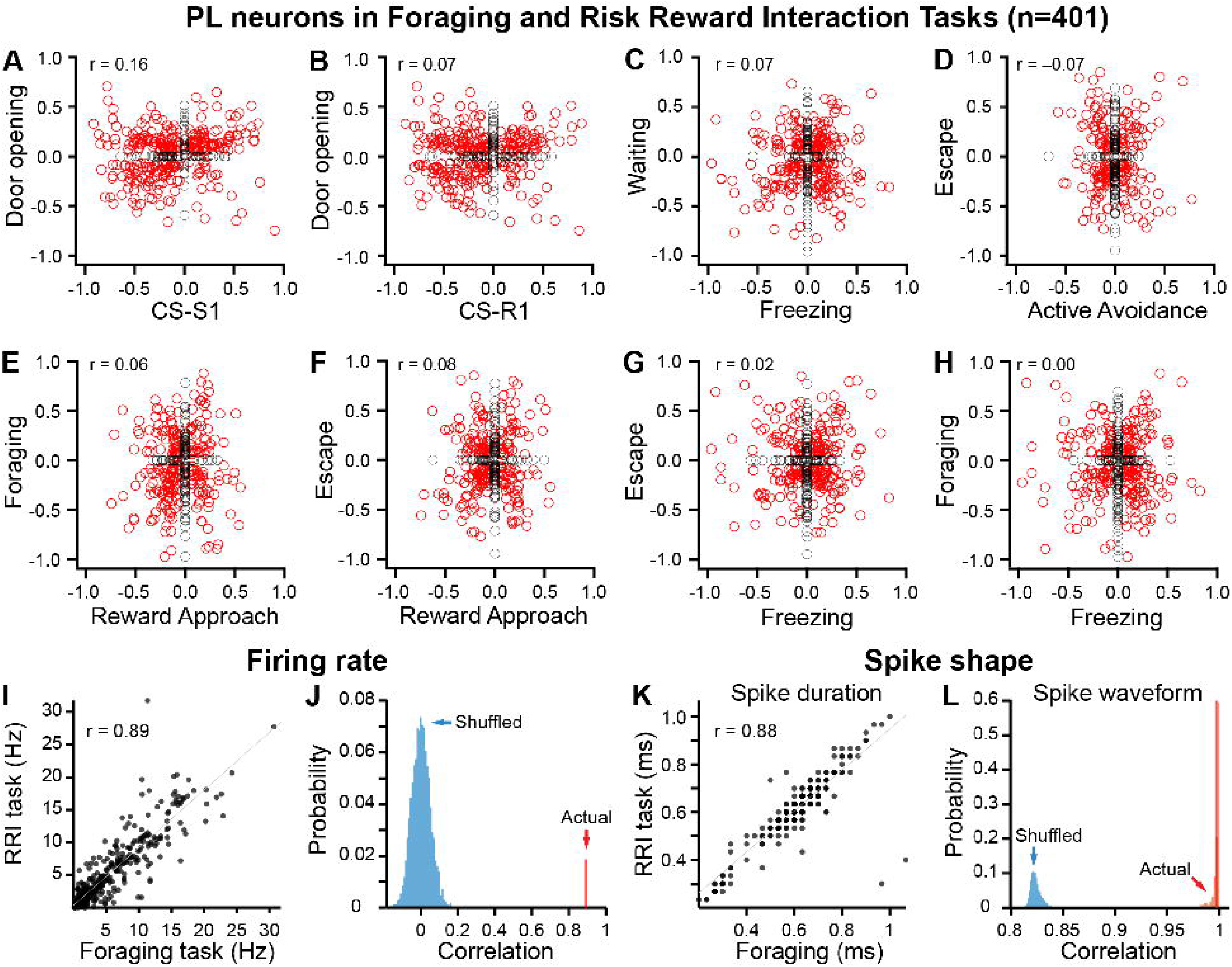
Correlation between the firing modulations associated with the variables of different tasks. Correlation between the GLM-estimated firing modulations associated with different variables of the RRI and foraging tasks. Each circle is a PL neuron. *Black circles:* neurons with zero modulation for one or both variables. *Red circles:* neurons with non-zero modulations. At the top left of each scatterplot is the Pearson correlation (*r*) between the variables. (**A,B**) Scatterplots relating the firing modulations linked to stimuli signaling the onset of trials. (**C-H**) Scatterplots relating the firing modulations linked to different behaviors. (**I**) Firing rates of the same cells during the RRI (y-axis) and foraging tasks (x-axis). (**J**) Comparison between the actual (red; *r* = 0.89) firing rate correlation between the two tasks to that obtained after shuffling cell identities in one of the two tasks with respect to the other 5000 times (blue; *r* = -3.4e-5±7.1e-4). (**K**) Spike duration of the same cells during the RRI (y-axis) and foraging tasks (x-axis). (**L**) Comparison between the actual (red; *r* = 0.997±2.9e-4) spike waveform correlations between the two tasks to that obtained after shuffling cell identities in one of the two tasks with respect to the other 5000 times (blue; *r* = 0.82±7.0e-5).

### Relation between multiple coding dimensions

Considering the ubiquitous two-way relationships found above, it follows that coding in cortical circuits is best described as a multidimensional lattice. Consistent with this, three-way scatterplots of the firing modulations form intricate but orderly eight-limbed X structures (**Fig. 6**). When rotating from a 2D to a 3D view, each of the X’s limbs can be seen to split in two along the z-axis, a phenomenon observed with most combinations of variables. See 3D animation (**Movie S1**) for a clearer depiction of these transformations.

**Figure 6.**
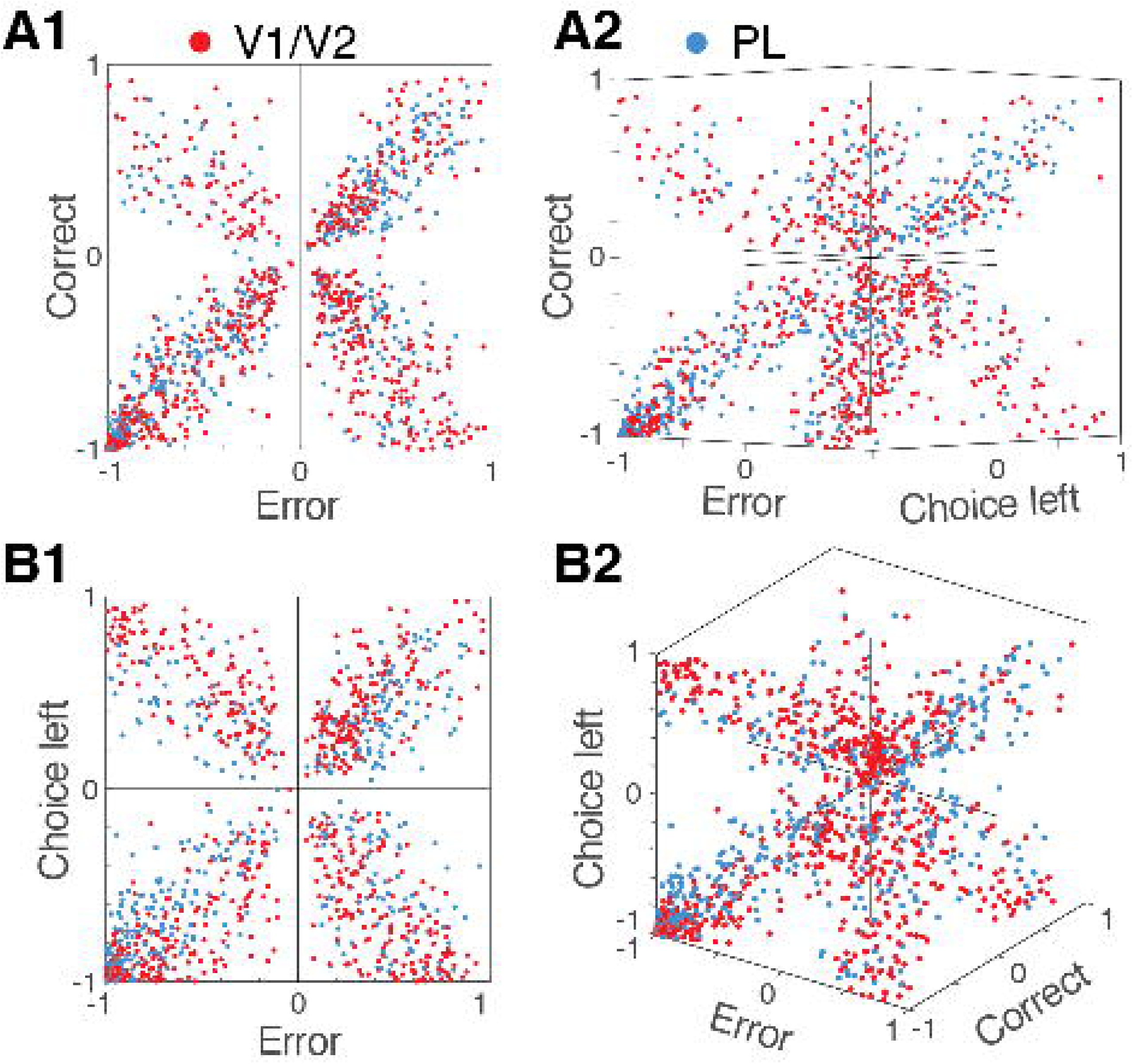
Relation between GLM-estimated firing modulations associated with three different variables of the operation task. Blue and red circles: prelimbic and V1/V2 neurons, respectively. 2D scatterplots (left) are rotated (right) revealing that neurons in each quadrant split in two groups when observed along a third axis. Related to **movie S1.**

To gain further insights into the coding structure, we applied uniform manifold approximation and projection (UMAP), a non-linear dimension reduction algorithm, to the modulations estimated in each task (**Fig. 7**). Specifically, we transposed the matrix of GLM-estimated firing modulations (*N* neurons X *D* variables) to embed the behavioral variables directly into a low-dimensional manifold (2D). Utilizing a correlation-based distance metric, UMAP positioned variables according to their population-level similarity without imposing linear constraints on the network geometry^32^. In the transposed state space, spatial proximity between variables signifies a shared neural representation; variables that map closer together recruit overlapping neuronal ensembles (**Fig. 7**).

**Figure 7.**
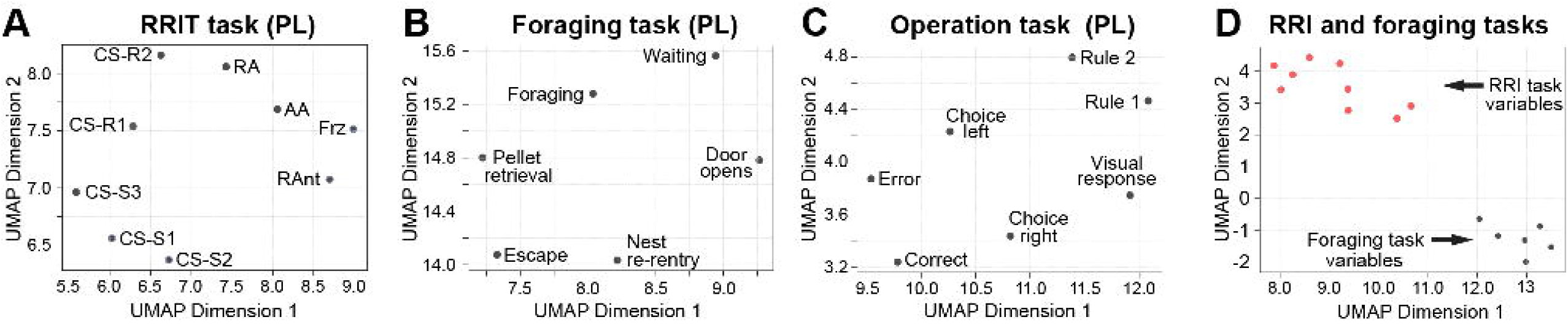
UMAP-based analyses of coding topology. Variable topology in the (**A**) RRI, (**B**) foraging, and (**C**) operation tasks in PL neurons (n=401). (**D**) Simultaneously clustered variables of the RRI (red circles) and foraging (black circles) tasks in neurons recorded during both tasks on the same day. Spatial proximity between variables indicates a shared neural representation.

Consistent with the positive correlations found between the firing modulations associated with the CSs in the RRI task, all five CSs were close together in the state space (**Fig. 7A**). Similarly, in the operation task (**Fig. 7C**), related variables like Rule 1 and 2, Correct and Error, or Left and Right lever were close to each other in the state space. By contrast, when we applied the same approach to the foraging and RRI modulations estimated in neurons recorded during the two tasks on the same day, the variables of the two tasks were completely segregated (**Fig. 7D**).

## DISCUSSION

Prior studies reported that neurons in higher-order brain regions simultaneously encode multiple task features^1–10^. However, neurons seemed to respond to random sets of task features^8,19,22^, often resulting in near zero correlations between the encoding of different variables at the population level^23^. While we too found a low correlation between the firing rate modulations associated with many of the variables, scatterplots revealed a highly structured organization consisting of correlated and anti-correlated axes, in line with the conclusions of a recent study on PL neurons in a single task^31^. Below, we highlight salient features of this coding structure and then consider its origin and implementation in different cortical areas.

### Coding structure in PL and V1/V2

The following applies to all types of variables except for those treated as arousal signals, a point we will return to. The defining feature of the coding structure we observed in PL and V1/V2 is that neurons with low or high absolute firing rate modulations by one variable tend to be modulated to a similar degree by other variables. However, the polarity of the modulations by one variable is not predictive of the modulations by another. For instance, when examining variables two at a time in a scatterplot, a sizable proportion of cells falls in all four quadrants (x^+^y^+^; x^−^y^+^; x^−^y^−^; x^+^y^−^; e.g. **Fig. 2**). Moreover, when the modulations of cells in one quadrant (x^+^y^+^ for instance) are plotted as a function of a third variable, many of them are seen to split further (z^−^x^+^y^+^ or z^+^x^+^y^+^ in this case; **Fig. 6A2**). Hence, the coding structure is not category-free in that response features are not randomly distributed across neurons but form functional clusters. Each additional variable potentially results in a doubling of the total number of categories. However, very large samples of neurons will be needed to examine this question as the proportion of neurons in each category decreases dramatically with the number of variables (and potential categories).

### Differences between the coding structure of PL and V1/V2 neurons

While it has been emphasized that neurons in higher-order cortical areas exhibit high-dimensional coding schemes, much evidence indicates that neurons in primary sensory regions also implement multidimensional coding^33–39^. Consistent with these findings, we found that the coding topology of PL and V1/V2 neurons was identical except for two aspects. First, modulations by visual stimuli were larger in V1/V2 than PL neurons. As a result, scatterplots relating modulations by visual stimuli to other variables had more similar adjacent angles in V1/V2 than PL neurons. Second, when relating modulations associated with the two rule visual cues, nearly all PL neurons fell on the main diagonal of the scatterplots. This contrasted with V1/V2 where a similar proportion of neurons fell in the two diagonals. We interpret these results as evidence that the modulation of V1/V2 activity by rule cues is dominated by their sensory features whereas PL neurons signal the presence of the trial phase. Thus, while the neocortex appears to rely on a uniform coding structure, its implementation is flexible enough to accommodate differences in the coding profile of each cortical region.

### Origin of the coding structure

We hypothesize that inhibitory interneurons play a critical role in shaping the coding topology. Specifically, we surmise that when driven to fire by extrinsic inputs, subsets of principal cells excite interneurons with similar modulations, generating feed-forward inhibition in principal cells with opposite modulations by one (x^+^ vs. x^−^) or more variables (x^+^y^−^ vs. x^−^y^+^). In turn, this would cause disinhibition of the first group of principal cells. These antagonistic cell groups likely emerge as a result of activity-dependent changes in the neurons’ extrinsic and intrinsic connectivity during training.

An interesting question in this context is whether the structured synaptic interactions described above shape the responsiveness of neurons across tasks. Consistent with earlier results^14,16–21^, when we related the modulations of neurons by variables of different tasks, we found little evidence of generalization across tasks. That is, the X-pattern was degraded as quantified by markedly reduced correlations between the absolute modulations in each quadrant by variables of different tasks compared to variables of the same task. This result suggests that the intricate synaptic architecture that underlies each task’s coding structure is only one instantiation of a flexible and scalable mechanism whose organization is adapted to the pattern of inputs associated with each task.

In PL neurons, we observed that the X-pattern was more pronounced in the operation task than the RRI and foraging tasks. At present, it is unclear whether this task-related difference in the expression of the coding structure is due to the unique properties of the tasks or the differing amount of training required for rats to master them. Specifically, the foraging task required next to no training, the RRI task about one week of training, and the operation task over four months of training. An important challenge for future studies will be to disentangle the influence of task properties vs. training regimens on the expression of the coding structure.

## Supporting information

Moview S1

## Resource availability

### Lead contact

Further information and requests for resources and reagents should be directed to and will be fulfilled by the Lead Contact, Denis Pare.

### Materials availability

This study did not generate new unique reagents.

### Data and code availability

- This study did not generate unique codes. The full dataset is available from the corresponding author upon request.
- This study did not report original code.

## Acknowledgements

This work was supported by R01 grant MH-130331 to D.P. from NIMH. M.B. was supported by the Graduate Program in Neuroscience of Rutgers University.

## Author contributions

M.B., A.A., I.T.K, D.B.H., and D.P. conceived the experimental design. M.B., A.A., I.T.K. and D.B.H. carried out the experiments. All authors contributed to analyzing the data. D.P. made the figures. D.P. wrote the first draft of the manuscript. All authors contributed to refine the manuscript.

## Declaration of interests

The authors declare no competing interests.

### Declaration of generative AI and AI-assisted technologies

The authors declare that they did not use generative AI or AI-assisted technologies when writing this paper.

## Supplemental information

Figures S1–S6 and Movie S1

## STAR METHODS

### Key resources table

| REAGENT or RESOURCE | SOURCE | IDENTIFIER |
| --- | --- | --- |
| **Antibodies** |  |  |
| None |  |  |
| **Bacterial and virus strains** |  |  |
| None |  |  |
| **Chemicals, peptides, and recombinant proteins** |  |  |
| DiI | Lumiprobe, Westminister, MD | Cat# 22366-93-4 |
| **Experimental models: Organisms/strains** |  |  |
| Long-Evans Hooded Rats | Charles River Laboratories | RRID: RGD_2308852 |
| **Software and algorithms** |  |  |
| MATLAB | MathWorks | RRID: SCR_001622 |
| Illustrator | Adobe | RRID: SCR_010279 |
| Igor Pro | Wavemetrics | SCR_000325 |
| Python 3 | Python™ | 3.10.0 |
| RStudio | https://rstudio.com | 2023.03.1 Build 446 |

### Experimental model and study participant details

#### Animals

We used Long-Evans rats of either sex that weighed 300-350 gm upon delivery (Charles River Laboratories, New Field, NJ; RRID: RGD_2308852). They were kept on a 12-hour light-dark cycle (lights-off at 7:00 PM). All procedures were carried out during the light phase of the cycle. All procedures were approved by the Institutional Animal Care and Use Committee of Rutgers University (PROTO201702617) and were in compliance with the Guide for the Care and Use of Laboratory Animals (DHHS).

### Method details

Fourteen rats of either sex (8 males and 6 females) were trained on one or two behavioral tasks and then implanted with Neuropixel probes in two or more of the following cortical regions: the prelimbic area (13 rats), posterior parietal cortex (5 rats), somatosensory cortex (8 rats), and visual cortex (5 rats). In the 12-14 days after delivery, rats had *ad libitum* access to food and water. During this period, they were habituated to the animal facility for one week and then to handling fifteen minutes daily for around five days. One week prior to the start of training, their access to food was restricted. The food restriction protocol consisted of restricting daily access to food in time while maintaining them at ≥85% of their initial body weight.

#### Risk reward interaction task

The behavioral apparatus was a dimly lit (10 Lux) rectangular arena (90 by 30 cm) with high polycarbonate walls and a floor made of evenly spaced metal bars (5 mm o.d., 8 mm spacing) which was divided in three sectors (each 30 by 30 cm). There was an array of light-emitting diodes (LEDs) behind the east and west walls as well as below each of the three floor sectors. At the east and west ends of the apparatus was a reward port. Using the LEDs, we delivered light conditioned stimuli (CSs; 20 s) that signaled different outcomes depending on their location. When presented underneath one of three shock sectors (CS-S), a footshock (0.4 mA, 10 s) was delivered in that sector, 10 s after CS onset. When presented behind the east or west walls (CS-R), a food reward was delivered at that location, 10 s after CS onset. The CS-Rs and CS-Ss were randomly ordered with 20-40 s inter-trial intervals. A white noise source was used to mask any sounds from the room during the task. A programmable microcontroller (Arduino, SparkFun, Niwot, CO) determined when and where the LEDs, shocks, and food rewards would be delivered.

Rats were first habituated to the arena for 1 hour. No light stimuli, food rewards, or shocks were delivered during this period, but the white noise maker was on. Over a period of 7-10 days, rats gradually learned to retrieve food rewards at the two ports and avoid shocks from each of the three sectors in response to the corresponding LEDs. On the first two days, rats were presented with 9 CS-S trials and 6 CS-R trials; once in the morning and once in the afternoon. Once rats learned to avoid the shocks, the sessions were extended to 39 CS-S trials and 26 CS-R trials. Training continued on days 3 through 7 with 1 one-hour session/day until rats reached at least 80% performance on all trial types combined. See^3^ for further details.

#### Foraging task

The behavioral apparatus was a rectangular enclosure (245 by 60 cm) with high polycarbonate walls. A retractable door (height, 50 cm; width, 10 cm) separated the apparatus into two sections: a dimly lit (10 Lux) nesting compartment (30 cm) with a water dispenser, and a longer (215 cm) and brighter (200 Lux) foraging arena. Trials began when the door to the nest was opened. On each trial, a food pellet (80-100 g) was placed at various distances from the nest (25-150 cm, in increments of 25 cm). The inter-trial interval lasted one minute. On a proportion of trials, a mechanical predator on wheels (Mindstorms, LEGO systems, Billund, Denmark; length, 34 cm; width, 17 cm; height, 14 cm) was placed at the end of the foraging arena, facing the nest. The predator had a sensor that detected the rats’ approach, causing a sudden advance (80 cm at 60 cm/sec) and jaw movements, after which it returned to its starting position. During recording sessions, rats were presented with alternating blocks of 10-20 predator and no-predator trials for a total of approximately 120 trials/day. During training, all trials occurred with no predator present. See^25^ for further details.

#### Operation task

Rats were presented with visual stimuli via two computer monitors placed 24 cm from their head (∼54 by 30.5 cm; 1366 by 768 pixels; 120 Hz refresh rate; ASUS TUF Gaming VG258QR, Taipei, Taiwan). Two epochs of visual stimulation occurred on each trial. (<u>1</u>) When rats stepped on a platform, they were shown one of two stimuli, identical on both sides, signaling one of two contexts (rules 1 and 2). These contextual stimuli indicated what dimension (speed or luminance) rats should ignore (distractor) and attend to (target) on a given trial. Under rule 1, rats had to attend to luminance and ignore speed. Under rule 2, they had to pay attention to speed and ignore luminance. (<u>2</u>) When rats nose-poked, different stimuli to be discriminated were presented on the left and right: moving filled circles on a black background. The circles varied along two dimensions: speed and luminance, both of which could assume five levels. Rats had to determine on which side (left or right) the target dimension was highest while ignoring the difference in distractor levels. Depending on trials, speed and luminance could be greater on the same or opposite sides (congruent and incongruent trials, respectively). Rats reported their decision by pressing a lever on the right or left, the side where the target dimension was higher. Upon pressing a lever, rats received immediate feedback on their performance. Correct responses caused a white LED to turn on above the reward port for 2 s in males and 1 s in females as rats were rewarded with food (Bio-Serv, Dustless Precision Pellets, 40 mg for males; 20 mg for females). Incorrect lever choices led the two screens to flash (four cycles of 0.25 s on, 0.25 s off). See^26^ for a detailed description of behavior in the operation task.

The psychophysics Matlab toolbox^40,41^ was used to control visual stimuli (https://github.com/Psychtoolbox-3/Psychtoolbox-3/tree/3.0.19.14). The stimuli to be discriminated consisted of 120 circles (40 pixels o.d.) of uniform luminance (9.53 to 100% illumination depending on the trial) that all moved downward at the same speed (30-1110 pixels/s) and occupied a random location within the screen. When a circle reached the bottom of the screen, a new circle appeared at the top at a random location. Stimuli started when rats nose-poked until they stepped off the platform or after one second, whichever came first. Luminance tests and adjustments were performed so that rats could detect increments in speed and luminance with equal ease (see^26^).

In each context, there were 500 possible combinations of target-distractor levels, (250 for trials where the target dimension was higher on the left or right). Depending on the target-distractor combination, trials varied in difficulty. Generally, the higher the left-right difference in target level, the easier the discrimination. When left-right differences in distractor levels pointed to the same side as the target’s, trials were easier than when they pointed to opposite sides. We did not present rats with trials with no left-right difference between the stimuli to be discriminated.

The various combinations of left-right differences in target and distractor levels defined 36 zones from which we selected trials. Prior to each session, a Matlab script (https://rutgers.box.com/v/AmirEtAlMatlabControlFiles) selected a sequence of left-right stimuli adapted to the rats’ proficiency in the prior session. Stimuli selection depended on the following sequence of control parameters: the chosen proportion of conflict trials, the probabilities associated with the possible differences in target (1-4) and distractor levels (0-4), and the number of trials devoted to each context. For each trial, the script determined from which of the 36 zones the left-right stimuli would be picked from, randomly selecting among the various possibilities that the zone in question contains. See^26^ for further details regarding the selection of stimuli to be discriminated, the training regimen, and performance in fully trained rats.

#### Surgery

Subjects were anesthetized with isoflurane (Henry Schein, Melville, NY) mixed with oxygen. After abolition of reflexes, they were placed in a stereotaxic apparatus (Kopf Instruments, Tujunga, CA) with non-puncture ear bars. Their temperature was kept at 37° C during the surgery and they were administered atropine sulfate (0.05 mg/kg, i.m.) to aid breathing. The scalp was shaven and then cleaned with Betadine and alcohol. Ten to fifteen minutes after injection of a local anesthetic along the scalp’s midline (Bupivacaine, 0.125% solution, s.c.), an incision was made, the skull was exposed and cleaned, and a burr hole was drilled above PL or the visual cortex. We used the following stereotaxic coordinates (in mm relative to bregma) from the stereotaxic atlas of Paxinos and Watson^42^ for PL (AP 3.5, ML 0.5, DV 3.2 to 4.8), V1/V2 (AP -7 to -8, ML 3 to 6, DV 0.5 to 3, with 10 degrees medio-lateral angle), S1 (AP -3.3 to -3.4, ML 5, DV 1 to 3.5) and the posterior parietal cortex (AP -3.8, ML 2.4, DV 0.4 to 2.5, with 10 degrees medio-lateral angle). Finally, a 3D printed headcap^43^ was cemented to the skull to protect the probe. Rats were allowed up at least one week to recover from the surgery.

#### Behavioral analyses

In the three tasks, behavior was monitored with an overhead video camera (The Imaging Source LLC, Charlotte, NC, USA) while neuronal activity was recorded extracellularly. Variations in position and running speed were determined using DeepLabCut^44^. We first obtained estimates of the rats’ position from videos. A residual Network (resNet) 101 from DeepLabCut’s “model zoo” was tuned by training it on 320 frames extracted from the videos of the rats performing the task, noting the rats’ head, shoulder, and tail. The videos were then processed by the tuned network to obtain position estimates at every frame, filtering and linearly interpolating for missing values. The rats’ position along the x and y axes were then processed in MATLAB (The MathWorks, Inc., Natick, Massachusetts, U.S.A) to score the rats’ behaviors.

In the RRI task, we considered the following four behaviors: reward approach, reward anticipation, behavioral freezing, and active avoidance. <u>Reward approach</u> started when rats began to move toward the reward port and ended when their nose reached the reward port. <u>Reward anticipation</u> started when rats placed their front paws on the reward port and waited there until reward delivery. <u>Freezing</u> was defined as periods of immobility (excluding breathing) that lasted at least one second during CS-S trials. <u>Active avoidance</u> began when rats first began to move off the lit sector ended when rats finished this behavior on one of the unlit sectors.

In the foraging task, using DeepLabCut pose estimations, we determined when rats started <u>waiting</u> at the door threshold (defined as when their snout extended beyond the door into the foraging arena), when they initiated <u>foraging</u> (defined as the last frame of immobility prior to fully moving out of the nest), retrieved of food pellet (hereafter termed “<u>food retrieval</u>”), escaped, and retreated into the nest. The observer also noted whether rats failed or succeeded each trial. The start of the <u>escape</u> phase was defined as when rats, after moving toward the food pellet, abruptly reverse direction and ran all the way to the nest. This behavior was observed irrespective of whether the predator was present or not and the trial was successful or not.

In the operation task, sensors detected when rats stepped on the platform, nose-poked, stepped off the platform, and pressed the left or right lever.

#### Histology

Localization of Neuropixel probes was assisted by coating their ventral extremity in a fluorescent dye DiI solution prior to implantation. At the experiments’ completion, rats were anesthetized with isoflurane and then perfused with a saline solution (0.9%) followed by a fixative (4% paraformaldehyde in 0.1 M phosphate buffer saline, Sigma Aldrich, St. Louis, MO). Their brains were then extracted, post-fixed for three to five days, and then immersed in a 30% sucrose solution for two days or more. Coronal sections (80 μm thickness) were obtained with a freezing microtome (Reichert, Depew, NY) and mounted on gelatin-coated slides to verify the position of the probe.

#### Recording and data processing

The data from the 384 channels of the Neuropixel probes was sampled at 30 kHz and recorded with a PXI-based acquisition card (IMEC, Belgium) controlled by the SpikeGLX software (https://github.com/billkarsh/SpikeGLX). The data was stored on hard drives. Recorded signals were processed offline using MATLAB.

Kilosort^45^ was used for clustering (https://github.com/MouseLand/Kilosort) and SortaSort (https://github.com/dbheadley/SortaSort) for manual refinement of the spike clusters identified by Kilosort. To distinguish spike clusters during manual refinement, we considered autocorrelograms and cross-correlograms. We required that autocorrelograms show a refractory period of ≥2 ms. Cross-correlograms could not have a refractory period, as this betrayed that the same unit was shared between clusters. We also inspected the data for spike shape reliability, spatially restricted spike waveforms, and continuous cell activity throughout the recording sessions. Spike trains were binned (50 ms).

### Quantification and statistical analysis

#### Assessing changes in firing rates in relation to task events and behaviors

To determine whether neurons showed significant changes in firing rates in relation to specific task events (stimuli or behaviors), we compared their firing rates before vs. during the event of interest using two-tailed signed-rank tests Bonferroni-corrected for the number of task events and behaviors considered in each task (9, 6, and 7 task events in the RRI, foraging and operation tasks, respectively). In the Results section, we report the number of cells that displayed significant changes in firing rates in relation to at least one event at p=0.05/n task events (0.0056, 0.0083, 0.0071 for the RRI, foraging, and operation tasks, respectively). See additional details below.

#### Computation of modulation indices

To examine task-related changes in firing rates on a common scale, we computed modulation indices whereby the firing rate change observed in response to a given task event was normalized to the neuron’s baseline firing rate. Slightly different approaches were used depending on the task based on their behavioral dynamics. We first describe the common points, then the differences, and conclude with various controls carried out to ascertain that our results were not dependent on the exact method used.

In the three tasks, we first measured the changes in firing rates associated with each task event by comparing each cell’s firing rate (averaged across trials) before the event onset vs. during the event. We then located the bin with the highest and lowest spike count during the event of interest, determined which had the highest absolute value relative to the pre-event baseline and accordingly classified the response as excitatory or inhibitory. Last, we normalized the cells’ maximal or minimal firing rate (FR_Peak_ or FR_Min_, respectively) for each task event to its pre-event baseline firing rate (FR_BL_) using the following equations:

For excitatory responses: (FR_Peak_ − FR_BL_) / (FR_Peak_ + FR_BL_)

For inhibitory responses: (FR_Min_ − FR_BL_) / (FR_Min_ + FR_BL_)

In the rare instances where the absolute difference between FR_Peak_ or FR_Min_ vs. FR_BL_ was equal, the cell’s modulation for this particular task event was given a value of zero.

The duration of the time windows used for the pre-event baseline varied between tasks based on the dynamics of behavior in each task: 5, 1.5, and 3 s for the RRI, foraging, and operation task, respectively. However, the duration of the time window for the event of interest was the same for all tasks (1 s). Also, when examining firing rate changes occurring in relation to behaviors of the RRI and foraging tasks, the last 750 ms prior to behavior onset were excluded from the pre-event baseline by imposing a –750 ms shift in the start of the baseline window. The rationale for this approach is our observation that firing rate changes related to behaviors consistently anticipated the onset of the behaviors.

Various controls were carried out to ensure that the X-pattern was not dependent on the exact method used to assess changes in firing rates associated with task events. A first alternative method consisted of determining the polarity of the response based on the difference between the average bin values of the pre-event vs. event periods but then using peak or trough value to quantify the modulation. Second, we compared the results obtained with the above two methods with vs. without normalizing the modulations by the pre-event baseline firing rate. All these approaches yielded qualitatively identical results in that the X-pattern was strongly expressed.

#### Thinning of spike trains

To test assess how neuronal firing rates influence the magnitude of the modulations, we excluded cells with firing rates <1.8 Hz. Then, cells with firing rates ≥1.8 Hz had their spike trains thinned to a maximum average firing rate of ∼2 Hz. To perform the thinning, we first computed the spike retention probability for each neuron as the ratio of the target rate to the observed rate. We then applied binomial thinning, where the number of trials is the observed spike count and the success probability is the retention probability. This produces a thinned spike train that is randomly subsampled from the original while preserving its temporal expectation structure.

#### Generalized linear models (GLMs)

Regularized regression, group Lasso GLMs, with Poisson distribution (grpreg R package^28^) and ten-fold cross validation was used to fit the spiking of individual cells for the duration of the tasks. Spiking was binned (50 ms bins) across the entire recording sessions. Variables were coded differently in the GLM depending on whether they were features that varied continuously or punctual events. Features that varied continuously like running speed assumed a range of values from 0 to 1. Punctual events were marked 1 when they occurred and zero otherwise. The stimuli and behavior events were convolved with basis functions defined by log-time scaled raised cosine bumps separated by π/2 radians (50 ms^46^). Each event kernel was represented as a linear combination of basis functions spanning a duration of time. Cross-validation sets were assigned by dividing each recording session into ten equal segments. This approach ensured that beta coefficients differed significantly from zero.

The GLM kernels for each stimulus and behavior that best fit spiking activity were generated by combining a set of pre-determined basis functions. Stimulus basis functions spanned their initial 10 s, had a rapid onset and narrow width which became more gradual and broader across time, reflecting stimulus responses that generally have sudden onsets and decay rapidly. By contrast, the basis functions used for behaviors started before and extended beyond the behavior onset to capture activity linked to planning and execution of the behavior. In both cases, these functions were adjusted by the GLMs’ beta values and summed within time bins to create a single kernel for each stimulus or behavior. If a unit did not encode a variable, the GLM gave beta values of 0 for the corresponding basis functions. Only parameters that best fit observed spiking are retained by the GLM, favoring dimensionality reduction.

The GLM of the RRI task considered the following nine main variables: CS-R1, CS-R2, CS-S1, CS-S2, CS-S3, reward approach, reward anticipation, freezing, and active avoidance. The GLM also considered three additional variables related to the rats’ position and movement speed but they are not considered here. The CS-Rs and CS-Ss were treated in the same way with basis functions covering their initial 10 s. They had a sharp onset and their duration increased with time from CS onset (Fig. S5A in^3^). Similarly, all the RRI behaviors were treated in the same way. They were described by a set 14 symmetrical basis functions (7 on each side) centered on the behavior onset and covering a total of 7.6 s. These basis functions progressively sharpened with time in the first 3.8 s and widened in the last 3.8 s (Fig. S5B in^3^).

The GLM of the foraging task considered the following seven main variables: door opening, waiting, foraging, pellet retrieval, escape, nest re-entry, and running speed. The GLM also considered an additional set of six variables (pellet position, predator presence, position, activation, aborted trial, running speed) but these are not considered here. Behaviors of the foraging task (waiting, foraging, escape, and nest-reentry) were coded in the same way. They were described by a symmetrical set of 20 basis functions centered on the behavior onset (10 on each side) and covering 4 s. They sharpened with time in the first 2 s and widened with time in the last 2 s.

In the operation task, we considered the following 7 variables: rule 1, rule 2, left lever, right lever, correct, error, visual stimulus. Running speed was also included in the GLM but is not considered here. For rule 1 and 2, the kernel spanned 3 s starting when rats stepped on the platform and was described by 10 basis functions whose duration widened as time passed. For choice left and choice right, the kernel spanned 1.5 s starting when rats nose-poked and was described by 10 basis functions whose duration widened as time passed. For correct and error, the kernel spanned 1.5 s starting when rats pressed a lever and was described by 10 basis functions whose duration widened as time passed. For the visual stimulus, the kernel spanned 1 sec starting when rats nose-poked and was described by 10 basis functions whose duration widened as time passed.

In each case, we estimated each variable’s contribution to the neurons’ firing rates and computed corresponding PETHs, aligned to the corresponding event. After calculating the PETH, the GLM-estimated firing rate (FR) modulation associated with each variable was determined using one of two equations:

For excitatory responses: (Peak FR – Baseline FR) / (Peak FR + Baseline FR)

For inhibitory responses: (Minimum FR – Baseline FR) / (Minimum FR + Baseline FR). Finally, the significance of the GLM-based modulation indices was assessed. The response value (peak or through) had to exceed ± 1.645 z of the baseline; otherwise it was set to zero.

#### Correlation between actual and GLM-predicted firing

To assess the goodness of fit of GLM estimates, for each task separately, we computed the Pearson correlation between the actual and GLM-predicted firing rate around all task events for each neuron. We then averaged the correlation coefficient across all neurons and variables. In the Results section, we report the grand average Pearson correlation±SEM for each task and the mode of the frequency distribution of the correlation coefficients.

#### Quantifying the properties of scatterplots

To characterize scatterplots relating GLM estimates of the firing modulations associated with different pairs of variables, the following properties were quantified. First, we computed the Pearson correlation between the firing modulations of all available cells, including cells with zero modulation for one or both variables. Second, we compared the ratio of neurons in the two diagonals as follows: n neurons in quadrants 2 and 4 / n neurons in quadrants 1 and 3. Third, we computed the standard deviation of the modulations along the x and y axes separately. Fourth, we computed the Pearson correlation between the firing modulations in each quadrant separately. When using this data for group comparisons, we first applied an absolute value function to the correlation coefficients.

#### Assessing whether the X-pattern can arise by chance

To test this possibility, for each task separately, we shuffled the GLM modulations of each neuron 1000 times. Each time, and for each pair of variables, we computed the correlations between the absolute modulations in the four quadrants separately. For each task, this resulted in an array of correlation coefficients whose length was equal to the number of variables pairs (which varied between tasks) times the number of quadrants (4) times the number of shuffles (1000). In each task separately, we then computed the actual average of the absolute correlations across all variable pairs and compared it to the null distribution.

#### Statistical analyses

All statistical analyses were conducted using MATLAB (MathWorks, Natick, MA) or Igor Pro (Wavemetrics, Portland, OR). Values are expressed as averages ± SEM. All statistical tests are two-sided and use a significance threshold of p = 0.05, Bonferroni-corrected for the number of comparisons. No data was excluded. We used Kruskal Wallis ANOVAs or Friedman tests when comparing three or more independent or related samples, respectively. Post hoc tests comparisons for significant effects varied depending on the type of data including Dunn, signed rank, or rank sum tests. In some cases, we generated null distributions by shuffling GLM-determined firing modulations 1000 times.

