## Supplemental figures 1-6 for "A UNIFORM CODING STRUCTURE IN THE NEOCORTEX"

SUPPLEMENTAL INFORMATION

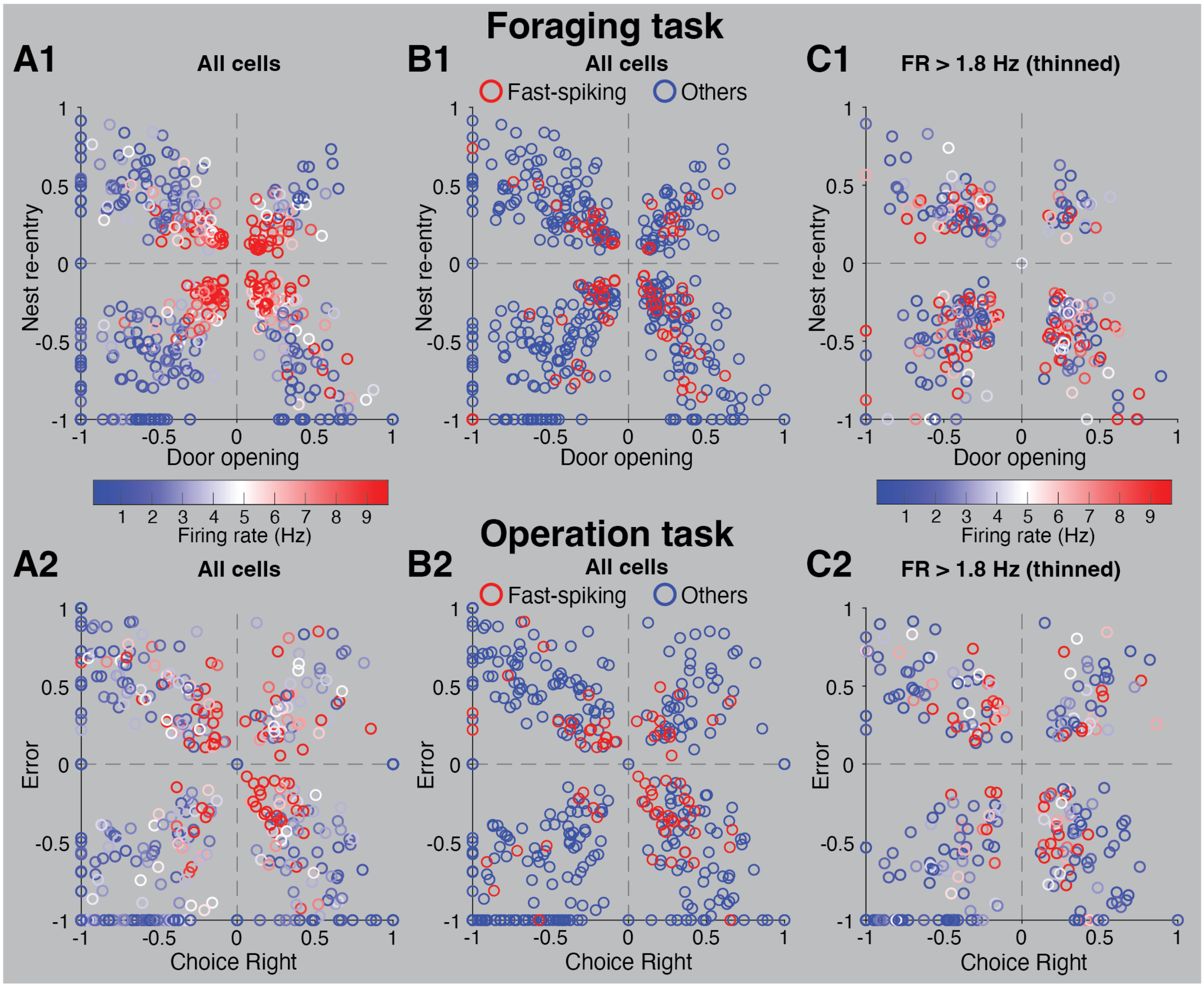


**Figure S1**. The X-pattern is not an artefact of the procedure used to normalize modulations by firing rates. (**A**) Firing rate modulations for two variables of the Foraging (**A1**) and Operation tasks (**A2**) color coded by firing rate. (**B**) Firing rate modulations for the same variables as in **A**, identifying presumed fast-spiking cells with spike durations <0.5 ms in red. (**C**) Firing rate modulations for the same variables as in **A** and **B**, considering only cells with firing rates (≥1.8 Hz) and after thinning their spike trains to achieve an average firing rate of around 2 Hz. Modulations are color coded by the original firing rates. Related to **Figure 2**.

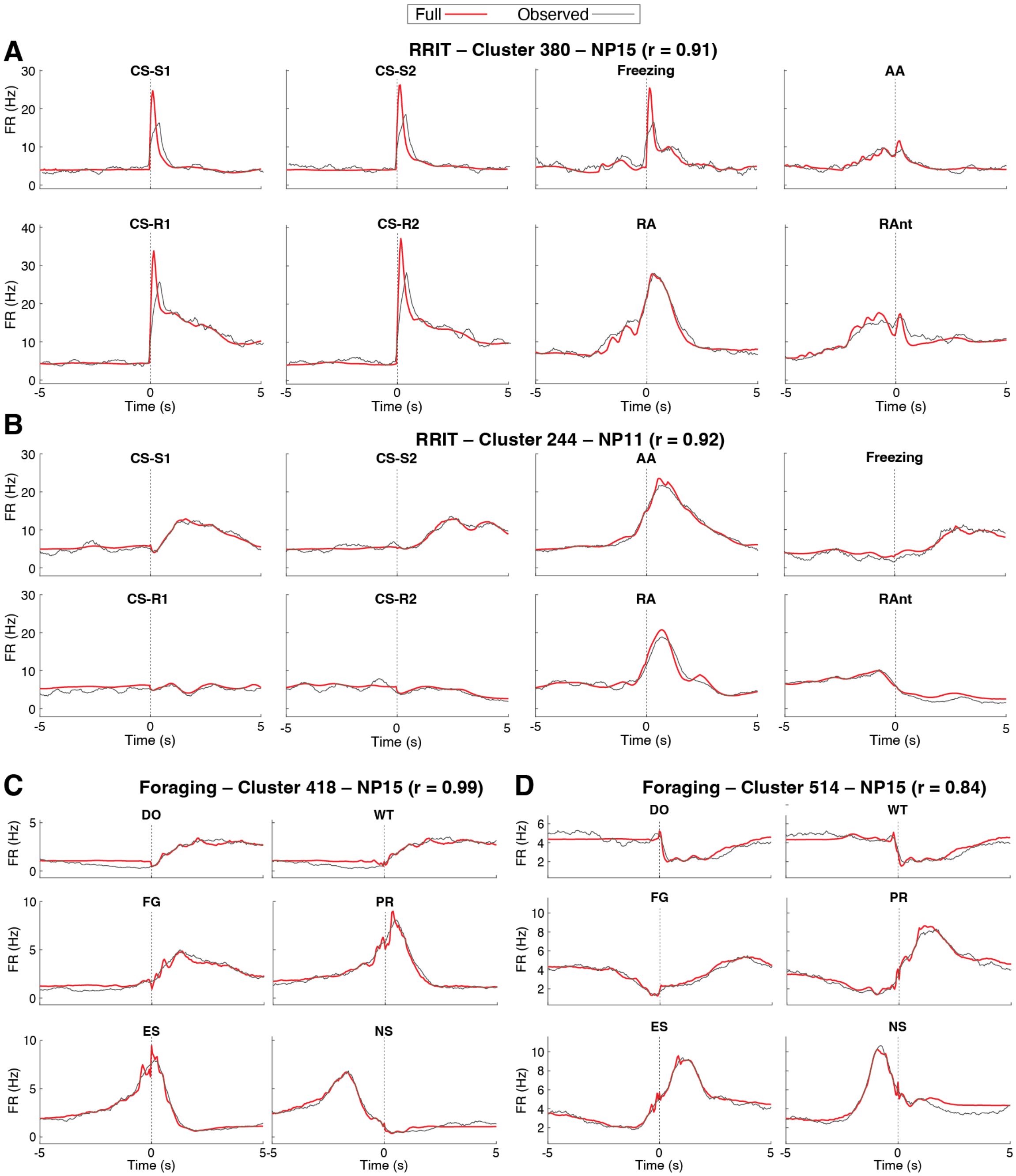


**Figure S2.** GLM-estimated coding in the RRI and foraging tasks. Coding of variables in the RRI (**A,B**) and foraging (**C,D**) tasks by four example PL neurons (**A-D**). Overlay of observed spiking (black lines) and estimated spiking (full model, red lines) for each condition (bold labels at top). Abbreviations: AA, active avoidance; DO, door opening; ES, escape; FG, foraging; NS, nest re-entry; PR, pellet retrieval; RA, reward approach; RAnt, reward anticipation; WT, waiting. Related to **figure 3.**

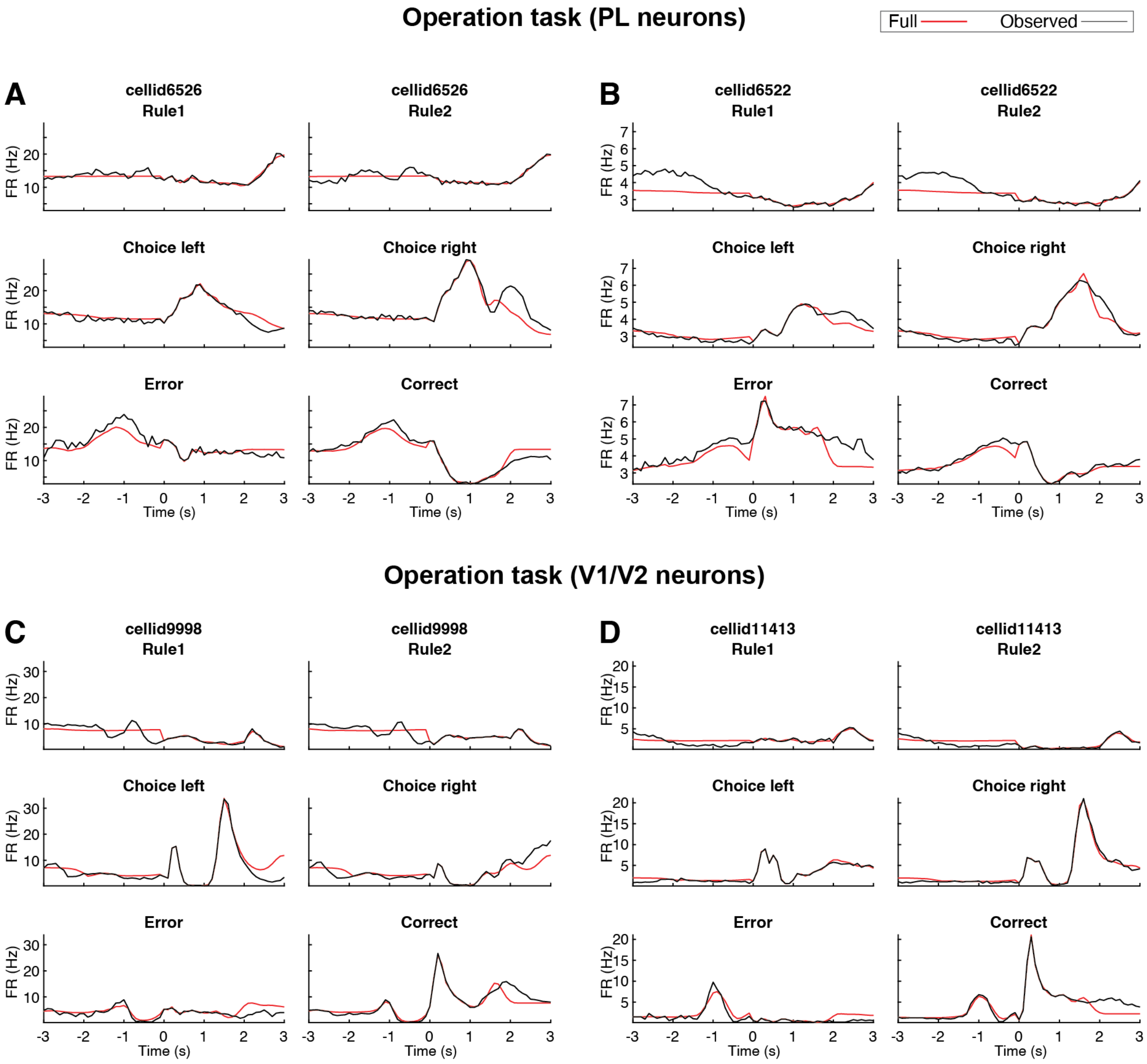


**Figure S3.** GLM-estimated coding in the operation task. Coding of variables in the operation task by two example PL (**A**,**B**) and V1/V2 (**C**,**D**) neurons. Overlay of observed spiking (black lines) and estimated spiking (full model, red lines) for each condition (bold labels at top). Related to **figure 3.**

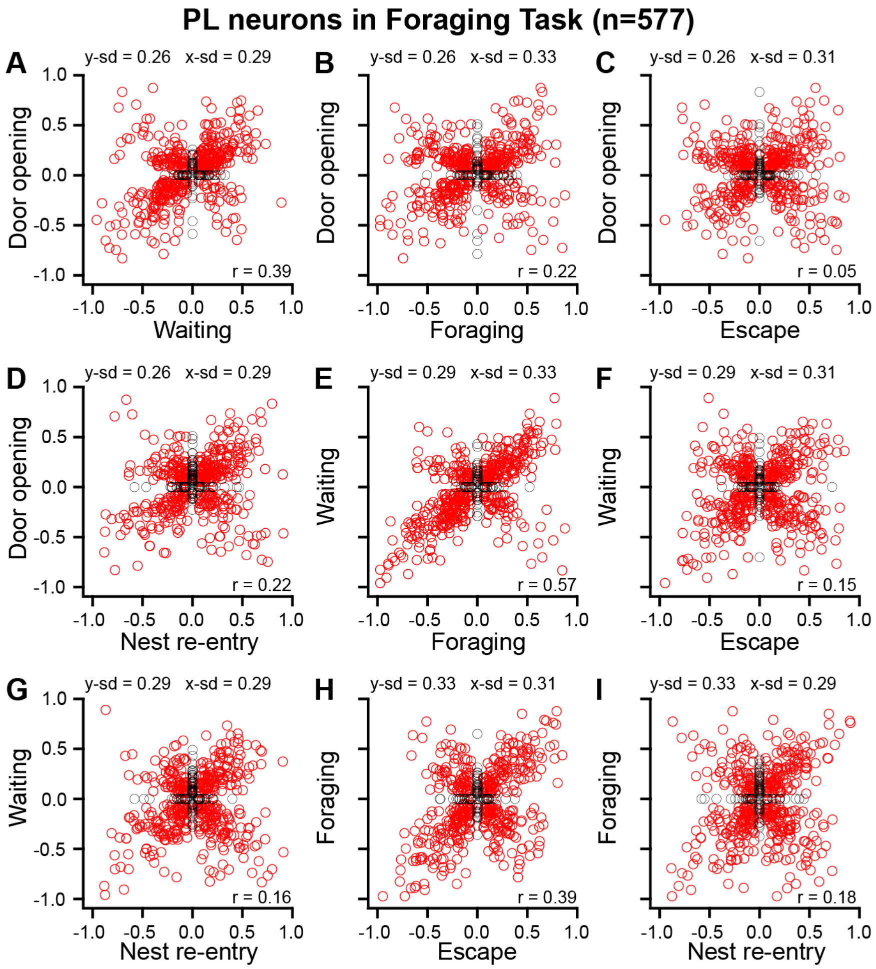


**Figure S4.** Coding structure in the foraging task. Correlation between the firing modulations associated with different variables of the foraging task. Each circle is a PL neuron. At the top of each scatterplot are the SD of the variable plotted long the y-axis (y-sd) and of the variable plotted along the x-axis (x-sd). The Pearson correlation between the variables (r) is listed on the lower right. Related to **figure 3.**

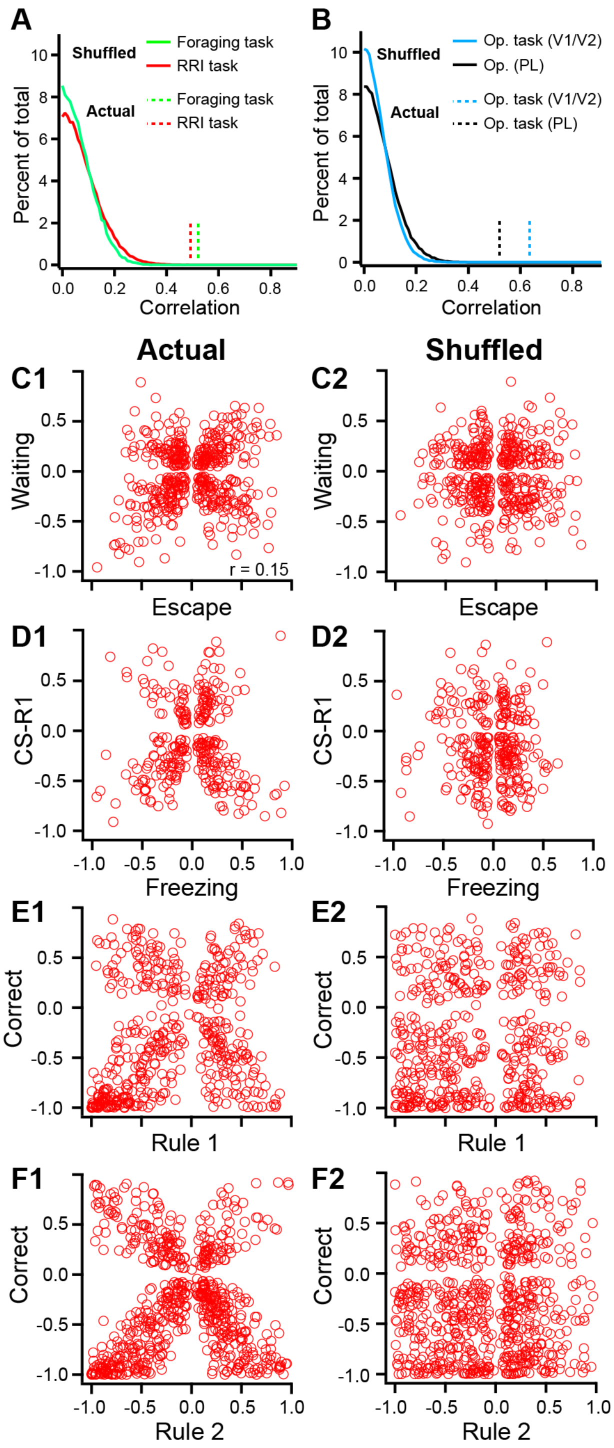


**Figure S5.** The X pattern is not a random occurrence. (**A,B**) Frequency distributions of correlation coefficients on randomized data in the foraging (green) and RRI (red) tasks (**A**) or the operation task (**B**) in PL (black) and V1/V2 (blue) neurons. To obtain these null distributions, each cell's modulations were shuffled 1000 times, each time computing the absolute correlations between the modulations in each quadrant for each pair of variables. (**C-F**) Scatterplots between actual (**panels 1**) and shuffled (**panels 2**) modulations (one example out of 1000) by variables of the foraging (**C**), RRI (**D**), and operation tasks (**E,F**) in PL (**C-E**) and V1/V2 neurons (**F**). For clarity, cells with zero firing modulations by one or both variables were removed. Related to **figure 3.**

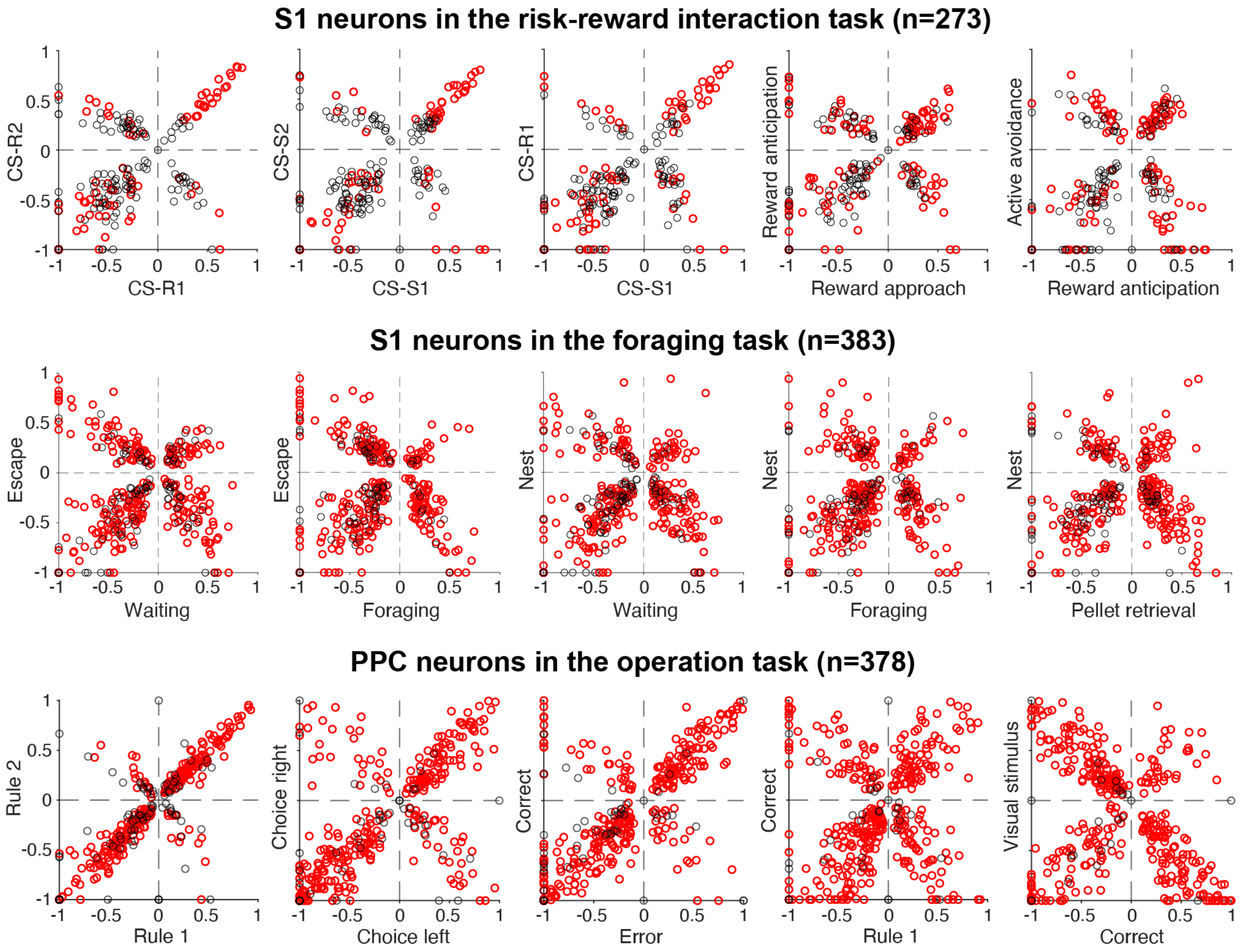


**Figure S6.** Correlation between the firing modulations associated with different variables of the RRI (top) and foraging (middle) tasks in S1 neurons and of the operation task in posterior parietal cortex (PPC) neurons. Each circle is a neuron. *Red circles*: neurons with significant changes in firing rates in relation to at least one of the task events or behaviors. *Black circles*: cells whose firing rate changes did not reach significance for either. Related to **figure 3.**
